# Cross-talk between Hippo and Wnt signalling pathways in intestinal crypts: insights from an agent-based model

**DOI:** 10.1101/492520

**Authors:** Daniel Ward, Alexander G. Fletcher, Martin Homer, Lucia Marucci

## Abstract

Intestinal crypts are responsible for the total cell renewal of the lining of the intestines; this turnover is governed by the interplay between signalling pathways and the cell cycle. The role of Wnt signalling in governing cell proliferation and differentiation in the intestinal crypt has been extensively studied, with increased signalling found towards the lower regions of the crypt. Recent studies have shown that the Wnt signalling gradient found within the crypt may arise as a result of division-based spreading from a Wnt ‘reservoir’ at the crypt base. The discovery of the Hippo pathway’s involvement in maintaining crypt homeostasis is more recent; a mechanistic understanding of Hippo pathway dynamics, and its possible cross-talk with the Wnt pathway, remains lacking. To explore how the interplay between these pathways may control crypt homeostasis, we extended an ordinary differential equation model of the Wnt signalling to include a phenomenological description of Hippo signalling in single cells, and then coupled it to a cell-based description of cell movement, proliferation and contact inhibition in agent-based simulations. Furthermore, we compared an imposed Wnt gradient with a division-based Wnt gradient model. Our results suggest that Hippo signalling affects the Wnt pathway by reducing the presence of free cytoplasmic β-catenin, causing cell cycle arrest. We also show that a division-based spreading of Wnt can form a Wnt gradient, resulting in proliferative dynamics comparable to imposed-gradient models. Finally, a simulated APC double mutant, with misregulated Wnt and Hippo signalling activity, is predicted to cause monoclonal conversion of the crypt.

## 1. Introduction

Colorectal cancer is the third most common malignant cancer, and the fourth leading cause of cancer death worldwide, accounting for roughly 1.4 million new cases and approximately 700,000 deaths in 2012 (1). Although the overall mortality rate for colorectal cancer has been declining by roughly 2% per year between 1997 and 2007 in the EU, from 19.7 to 17.4/100,000 men and from 12.5 to 10.5/100,000 women (2), mainly due to improved early diagnosis and/or an improved lifestyle, colorectal cancer still affects a significant number of people worldwide. Improvement of these statistics (through, for example, new diagnostics and treatment) will depend on a better understanding of the cellular mechanisms that are responsible for the disease. It has been suggested that there are specific cancer stem cells that develop and propagate colorectal cancer (3). Within the intestine, stem cells reside within crypts, tubular indentations lining the walls of the gastrointestinal tract (4) that are responsible for the total cell renewal of the intestine lining every 4-7 days in humans (5–7) (more rapidly in mice (8)). In a healthy crypt, regulatory networks tightly control individual cell and crypt homeostasis through multi-level processes such as proliferation, cell adhesion, growth factors and production of other secreted molecules. Changes to these underlying regulatory networks due to mutations can disrupt crypt dynamics and eventually result in tumorigenesis. Of particular significance are the Wingless/Int (Wnt) and Hippo signalling pathways.

The role of the canonical Wnt signalling pathway (i.e. the Wnt/β-catenin pathway, hereinafter referred to as Wnt pathway) in regulating the expression of target genes linked to proliferation and differentiation has been extensively studied, both experimentally and theoretically (9–12). Within the gastrointestinal tract, it is known that Wnt signalling governs proliferative dynamics specific to the lower regions of intestinal crypts (13). The importance of the Wnt pathway in maintaining a healthy cell turnover is clear from studies reporting consequences of Wnt signalling dysfunction (13–16), with the development of intestinal tumours and polyps linked to mutations in the signalling pathway cascade, and approximately 80% of all human colon tumours reported to show mutations that inactivate the APC gene (17). The main cellular effect of activation of the Wnt signalling pathway is to alter the subcellular localisation of β-catenin (9, 15). In the absence of a Wnt signal (‘Wnt-Off’), cytoplasmic β-catenin is rapidly degraded by the destruction complex formed by APC, Axin, Ser/Thr kinases GSK-3 and CK1 (9). In the presence of a Wnt signal (‘Wnt-On’), extracellular Wnt ligands bind to Frizzled cell surface receptors, causing inhibition of the destruction complex via sequestration of β-catenin. This leads to the accumulation of unphosphorylated β-catenin in the cytoplasm and thus increased shuttling of β-catenin to the nucleus, where it binds to TCF/LEF transcription factors and activates expression of Wnt target genes, many of which regulate cell proliferative dynamics (10). Typically, malignant transformations of cells within the crypt involve mutations in the APC gene (18), causing aberrant build-up of β-catenin and expression of Wnt target genes (10, 11, 14, 19). Cell proliferation occurs predominantly in the bottom third of the crypt (20), with a low level of extracellular Wnt detected at the crypt orifice and a maximal level at the base (14, 15, 21). Previous studies have suggested that such a Wnt gradient helps to maintain crypt homeostasis (specifically, constancy in the numbers of different cell types, and overall crypt size), determines subcellular β-catenin localisation and kinetics and, ultimately, regulates cell-cycle dynamics (22, 23). Indeed, many theoretical models of crypt dynamics have linked an imposed Wnt gradient directly to cell-cycle duration (5, 24–26). However, more recent experimental data suggest that the Wnt gradient results from a cellular division-based spreading of Wnt from the base of the crypt, where there is an effective reservoir of Wnt, and consequent Wnt dilution throughout the crypt (27).

Aside from Wnt signalling, other processes and signalling pathways also play important roles in crypt homeostasis. For example, contact inhibition (CI) of cell proliferation is defined as the cessation of cell-cycle progression due to contact with other cells, leading to a transition to a dense monolayer of epithelial cells (28). Increased CI can be due to forced reduction in cell volume, externally applied stress, or increase in the cell density, with solid stress shown to inhibit the growth of cell spheroids *in vitro*, regardless of differentiation state (29). Overcrowding can also induce live cell extrusion from the monolayer (30); however, with regards to crypt homeostasis, this appears to occur towards the crypt orifice, thus beyond the region we would expect proliferative cells to reside. We consider CI in our investigation, as the precise nature of the mechanisms driving it, possibly including the interplay of signalling pathways regulating crypt homeostasis, are not well understood (31, 32).

It has been suggested that the Hippo signalling pathway, which is deregulated in multiple cancers (33, 34), plays a role in preventing cell proliferation due to CI (35), with signalling linked to the volume of a cell and the overall stress applied to it (35–37). The Hippo pathway negatively regulates Yes-associated protein (YAP) and the transcriptional co-activator PDZ-binding motif (TAZ); the activation of YAP and TAZ promotes cell proliferation and inhibits cell death (28, 32). The Hippo pathway restricts the availability and functionality of YAP in the nucleus by altering its level and distribution (31, 38). Over-expression of YAP or its over-activation by Hippo pathway mutations have been shown to counter the effects of CI *in vitro* and organ size control *in vivo*, promoting tissue overgrowth and cancer development (32, 33). The Hippo signalling pathway, unlike other signalling cascades, does not appear to have its own dedicated extracellular peptides and receptors, but instead relies on regulation by a network of upstream components and mechanisms, such as cell polarity complexes and adherens junctions (39). It has been suggested that the Hippo pathway is regulated by the cellular architecture and the mechanical properties of cell environment, possibly serving as a sensor for tissue structure and mechanical tension (36, 40, 41).

The cross-talk between the Hippo and Wnt signalling pathways has been shown to play a key role in mediating cell proliferation (35), by reducing nuclear β-catenin levels and, consequently, the expression of Wnt target genes that control cell-cycle progression. In a Hippo ‘off’ state, the level of YAP-P remains stable within the cell, and the YAP-P/β-catenin complex remains at minimal levels in the cytoplasm. Conversely, in a Hippo ‘on’ state there is an increase in YAP-P, which binds to free β-catenin in the cytoplasm creating a YAP-P/β-catenin complex that is unable to localise to the nucleus of the cell, whilst maintaining its membrane bound activities. These results motivate our development of a mathematical model able to capture the mechanisms linking Wnt and Hippo signalling and CI, extending a previously proposed kinetic model of β-catenin within an individual cell (42).

Mathematical modelling has been used to explore crypt dynamics in health and disease, and to better understand how colorectal cancer can emerge from the interactions across several scales of biological processes, such as mutations that cause an abnormal intracellular response to changes at either the cell or tissue levels, and cell proliferation, motility and adhesion (43). Existing mathematical models of crypt dynamics have focussed on Wnt signalling, including its role in monoclonal conversion (the process by which a crypt is populated in its entirety by the progeny of a single ancestor cell (24, 44)), and the effects on subcellular β-catenin localisation on the cell cycle (25). Recently, the possible effects of cell-cycle cessation caused by contact-dependent factors have been investigated by modelling CI as cell-cycle cessation linked to cell volume (5). In all these models, Wnt is assumed to act on cells via an imposed and external gradient. Earlier models assumed that Wnt affects cell-cycle progression dynamically, while a more recent work suggests that cell-cycle duration is dependent on the Wnt signal available immediately after mitosis (5).

In what follows, we consider two Wnt models in the multicellular context: the first assumes that Wnt exists as an externally-imposed gradient, with variants as described above, where the Wnt level received by each cell either changes throughout its cell cycle or is updated at the point of each cell division. The second model considers the Wnt level to be an intrinsic property of each individual cell, instead of being prescribed only externally, and hence allows us to test the hypothesis of an emergent Wnt gradient in the crypt. In this second model, Wnt is distributed between the daughter cells following each cell division, with a Wnt source ‘reservoir’ region located at the base of the crypt. In the case of the small intestine this is akin to the region of Paneth cells that can transfer Wnt to neighbouring cells, although this diffusion has been shown to be limited to 1-2 neighbour cells (27, 45). A more recent study has suggested an additional source of Wnt proteins in FOXL1+ telocytes, which form a subepithelial plexus extending from the stomach to the colon, again localised towards the base of the crypt (46).

More fundamentally, models of crypt dynamics that explicitly include the Hippo signalling pathway are lacking, preventing a thorough analysis of the effects of Wnt/Hippo crosstalk. To address this, we propose a new model that describes both Hippo and Wnt signalling pathway dynamics, and links their activity to cellular proliferation and CI. We initially consider the effects of Hippo and Wnt at the single-cell level, and analyse their combined role in cell-cycle cessation. Moving to a multicellular framework, we show that crypt dynamics remain fundamentally unaltered upon incorporation of CI via Hippo signalling; the latter, however, plays a crucial role in the monoclonal conversion probabilities upon mutations in Wnt pathway genes. Whilst the mutation in Wnt pathway genes effectively cause increased proliferation in mutant cells throughout the crypt, the addition of Hippo signalling mediated CI causes a reduction in the proliferation of the healthy cells during the mutant monoclonal conversion process.

## 2. Computational Methods

The single-cell model combines the subcellular Wnt/Hippo signalling model, as described by a set of 9 Ordinary Differential Equations (ODEs, Figure S1), and the cell-cycle model through 5 additional ODEs (Figure S2). The Wnt/Hippo signalling model adapts a formalism previously proposed (25, 42) to describe the dynamics of β-catenin and the effects of signalling (Wnt) on the cell-cycle, to further include Hippo-signalling-dependent complex (β-catenin/YAP-P, Equation 5, Figure S1) and its effect on free β-catenin. The cell-cycle model used (both equations and parameters) is as in Swat *et. al.* (22). The agent-based model was implemented within the Chaste (v. 3.3) modelling framework (52, 53). Further details about model assumptions, equations, parameter values and simulation settings can be found in Supplementary Information.

## 3. Results

### 3.1 Intracellular modelling of Wnt/β-catenin and Hippo signalling pathways

We developed an ordinary differential equation (ODE)-based kinetic model to capture the combined effects of Wnt/β-catenin and Hippo signalling on intracellular dynamics. Our model builds on the formalism proposed by van Leeuwen *et al.* (42), which distinguishes adhesive and transcriptional functions of β-catenin. Although alternative models of Wnt signalling exist (47–50), the van Leeuwen formalism was chosen as it incorporates important mechanistic features of the canonical Wnt pathway (including sequestration of β-catenin by the destruction complex, and activation/inactivation of the destruction complex) and couples β-catenin localisation with cell-cycle progression (22). Specifically, when β-catenin localises at the cell membrane, it regulates the formation of E-cadherin-dependent cell-cell contacts, connecting adherens junction proteins to the actin cytoskeleton (42). Accumulation of β-catenin within the cytosol results in its nuclear translocation, activating the transcription of target genes linked to cell-cycle progression and cell-survival. Thus, cytoplasmic β-catenin can either: (a) be sequestered by molecules linked to Wnt gene transcription to form transcriptional complexes, (b) form adhesive complexes at adherens junctions, or (c) undergo Wnt-mediated degradation.

We extended the Wnt model to include a phenomenological description of the effect of Hippo on Wnt signalling, and to implement cell volume-dependent CI. As postulated by Imajo *et al.* (35), we modelled the Hippo signalling-governed level of cytoplasmic phosphorylated YAP to cause a reduction in the levels of nuclear β-catenin. This is in line with experiments showing that Hippo signalling causes increased phosphorylation of YAP, combined with the formation of YAP-P/β-catenin complexes within the cell but constrained to the cytoplasm (31, 34, 35). We assumed that the mechanism of CI is a reduction of cell volume below the equilibrium volume of a typical crypt epithelial cell, as this should correlate with an increase in surface stress due to cellular crowding in an epithelial monolayer, as well as an increase in cell density (29). We therefore altered the kinetic model (Figure S1) to include volume-dependent nuclear sequestration rate of β-catenin (35); in such a way, volume-dependent Hippo signalling can decrease nuclear accumulation of β-catenin (thus increasing its cytoplasmic presence without altering its total cellular level) and, consequently, alter cell proliferation dynamics (Supplementary Information).

Figure 1a shows schematically the interplay between Wnt and Hippo signalling within a single cell in our model. This is the basis for the kinetic diagram in Figure 1b, representing interactions between free Axin (*X*), adhesive molecules (*A*), the active destruction complex (*D*), the β-catenin/YAP-P complex (*C*_*H*_), four molecular forms of β-catenin (*C*_*i*_, *i* = *A*, *C*, *T* and *U*, corresponding to β-catenin contained in the adhesive junction, cytoplasm, transcriptional complexes, and marked for ubiquitination/degradation, respectively), transcriptional molecules (*T*), and the Wnt target protein (*Y*).

**Figure 1.**
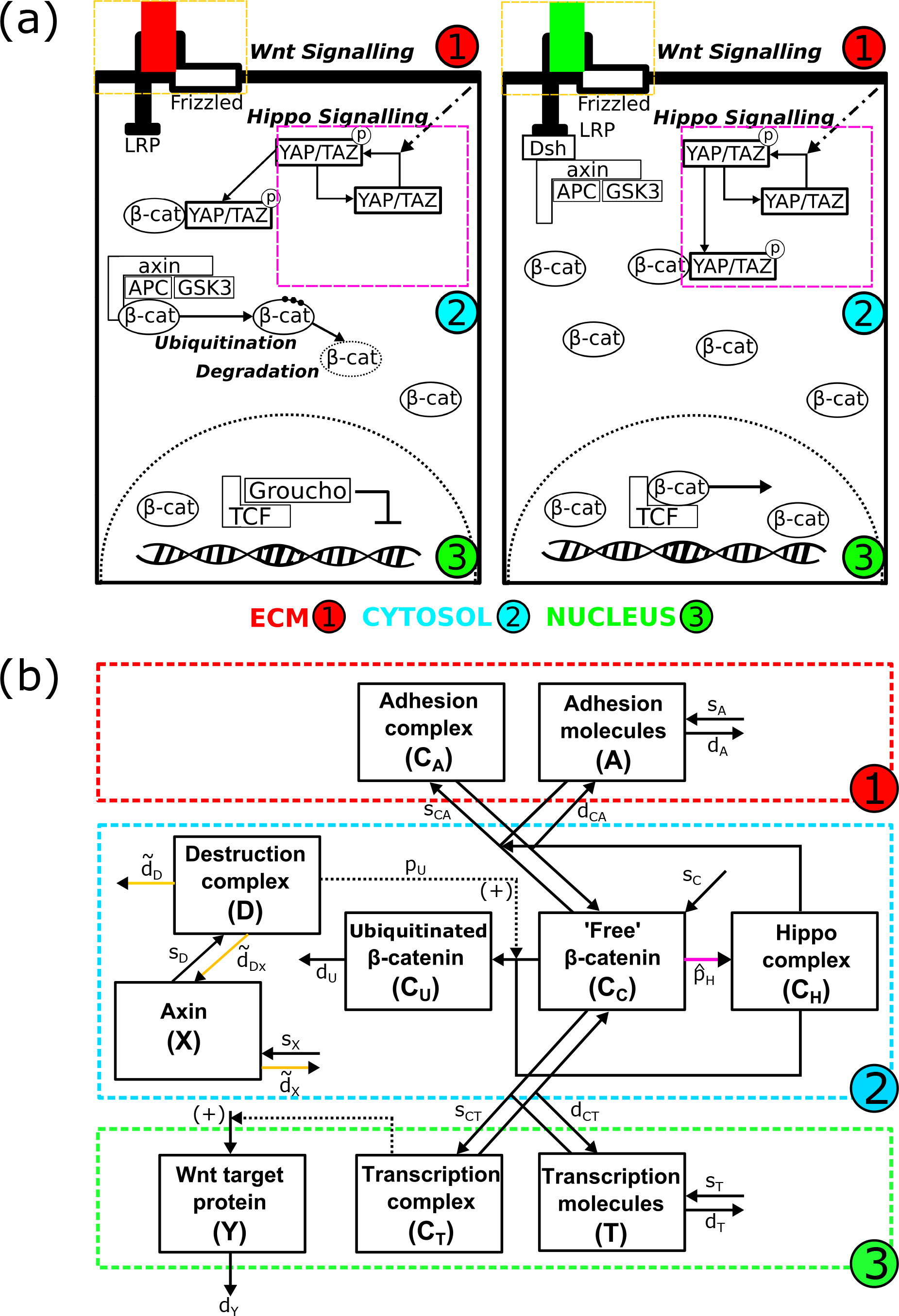
Wnt/Hippo signalling network. **(a)** Schematic showing β-catenin cellular localisation, dependent on Wnt and Hippo signalling. β-catenin exists either in the cytosol or in the nucleus, with its localisation directly affecting cell-cycle progression. In the Wnt-Off state (left panel), β-catenin is degraded in the cytosol by a complex comprised of APC, Axin and GSK3, preventing β-catenin nuclear localisation. In the Wnt-On state (right panel), the destruction complex is disrupted by Disheveled (Dsh), resulting in increased β-catenin levels and its nuclear localisation. This causes the displacement of Groucho in the nucleus and transcription of Wnt target genes linked to proliferation and cell-cycle progression. Hippo signalling causes the phosphorylation of YAP/TAZ within the cytosol. Phosphorylated YAP/TAZ binds to cytosolic β-catenin, preventing its nuclear accumulation and, in turn, transcription of Wnt target genes and cell-cycle progression. **(b)** Network diagram resulting from the schematic in (a) and describing the kinetics within the cell. *C*_*A*_, *C*_*C*_, *C*_*T*_, and *C*_*U*_ are the levels of adhesive-linked β-catenin at the cell surface, cytosolic β-catenin, transcriptional nuclear β-catenin, and β-catenin marked for degradation, respectively. *A*, *T* and *D* denote the level of molecular species forming complexes with β-catenin at the cell surface (forming adhesive complexes at the adherens junction), within the nucleus, and within the destruction complex, respectively. *X* and *Y* denote the levels of Axin and transcribed Wnt target proteins, respectively. *C*_*H*_ denotes the level of β-catenin/YAP complex formed due to Hippo signalling in the cell. Rates which depend on activity of signalling pathways are indicated by the coloured arrows (yellow for Wnt dependence, pink for Hippo dependence).

The temporal Wnt/Hippo dynamics in a single cell were modelled as a system of Ordinary Differential Equations (ODEs, Figure S1); they augment a previously derived and fitted model of Wnt pathway in intestinal cells (25) to phenomenologically describe both the aforementioned effects of Hippo signalling through the phosphorylated YAP/β-catenin complex, and CI (details in Supplementary Information). The Wnt/Hippo model is mainly based on mass-action and Michealis-Menten kinetics (51); it was derived from the original publication (25), and, as such, the parameters were left unchanged where used previously (Figure S1, Table S1). As an extension to the original model, we described the dynamics of the β-catenin/YAP-P complex (*C*_*H*_, Equation 5, Figure S1) to account for the effects of Hippo signalling on β-catenin localisation (Figure S1). As in (25), we also modelled the interplay between the pathway signalling and the cell-cycle; this is governed by the amount of transcriptional β-catenin (*C*_*T*_), which couples the signalling ODE model (Figure S1) to the cell-cycle model (ODEs in Figure S2, equations and parameters unchanged from the original work in (25); further details in Supplementary Information).

To explore the effect of Hippo signalling on cell-cycle duration and fix the parameters in the Wnt/Hippo model not present in the original model (i.e. those in Equation 5, Figure S1), we performed a parameter sensitivity analysis, varying the YAP-P/β-catenin complex binding rate (*p*_*H*_) and dissociation constant (*K*_*H*_); these parameters govern the effectiveness of the inhibition of nuclear β-catenin localisation and, consequently, CI. Of note, in the single-cell model (Figure S1), *p*_*H*_ is varied within a given range (i.e. takes specific values as cell volume is not directly modelled), while, in the agent-based model (see Section 3.2), 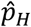 is dependent on cell volume (i.e. 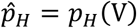, further details in Supplementary Information). Single-cell simulation results, shown in Figure 2a, suggest that the saturation coefficient (*K*_*H*_) has some effect on cell-cycle duration but, more significantly, alters the effective range over which the binding rate (*p*_*H*_) controls cell-cycle duration, with changes in the binding rate having the largest effect on delaying and eventually stopping proliferation. We therefore assigned a value to the saturation coefficient *K*_*H*_ (Table S1) which, when combined with Wnt signalling effects on β-catenin localisation, did not cause premature cessation of the cell cycle across the required range of Wnt signal.

**Figure 2.**
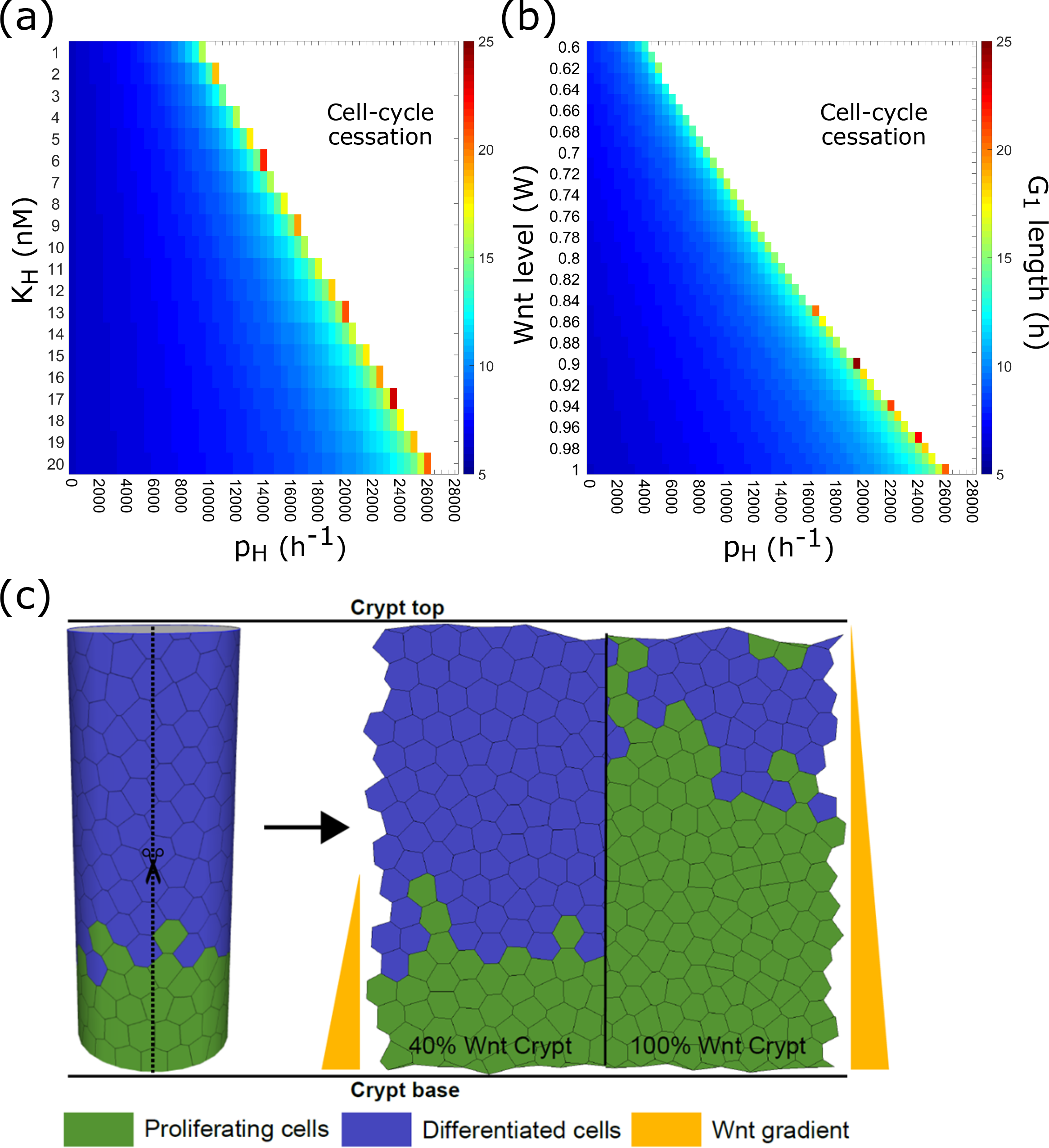
Single-cell ODE model of Hippo/Wnt signalling network. **(a)** Single-cell sensitivity analysis of the Hippo signalling module; parameters *p*_*H*_ and K_H_ represent the binding and dissociation rates of phosphorylated YAP/β-catenin and its complex, respectively. The coloured region shows the length of the cell cycle, and the white region represents cell-cycle cessation. **(b)** Single-cell sensitivity analysis of the Hippo-dependent cell cycle as a function of (static input) Wnt level and *p*_*H*_, for *K*_*H*_ = 20nM. The Wnt signal affects the β-catenin destruction complex (Equations 1 and 2 in Figure S1) according to relationship in Figure S1 legend. **(c)** Schematic diagram of the 3D to 2D projection method, ‘unrolling’ the cylindrical crypt to a planar domain with periodic boundary conditions. The right panel illustrates the imposed-gradient Wnt model M_E_, in the two considered cases of minimum Wnt level at 40% and 100% of the height of the crypt, and the resultant effect on cell proliferation region.

We then investigated how the cross-talk between the Hippo and Wnt pathways affects the duration of the cell cycle (*i.e* the time taken for the levels of *E2F1* gene -Equation 2, Figure S2- to surpass the threshold *E2F1* > 1 for initiating the transition from the G_1_ to the S phase of the cell cycle, as in Swat *et al.* (22)). In single cell simulations, the Wnt signalling level (*W*) was indirectly modelled through Wnt-dependent parameters affecting the β-catenin destruction complex (see Equations 1 and 2 in Figure S1, where the parameters carrying tildes 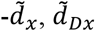 and 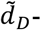 vary in response to Wnt signalling, as described in Supplementary Information). Setting the saturation term *K*_*H*_ to 20nM to allow for an appropriately sized range over which to vary the binding rate, a concurrent increase in the YAP-P/β-catenin complex binding rate (*p*_*H*_) and decrease in Wnt signalling level caused a delay and eventual cessation of the cell cycle (Figure 2b). Whilst the choice of *K*_*H*_ = 20 nM and *p*_*H*_ = 26000 h^−1^ are not unique, they are chosen to prevent premature cell-cycle cessation at lower levels of Wnt. An alternative choice of *K*_*H*_ and *p*_*H*_ would still cause cell-cycle cessation in response to volume change; however, smaller values of *K*_*H*_ would result in more rapid cell-cycle cessation in cells experiencing lower Wnt signalling than we would expect.

These results suggest that cells experiencing low Wnt signalling levels are more susceptible to inhibition of proliferation by Hippo signalling, as there is less free β-catenin localised in the cytoplasm. This single-cell analysis also shows that the reduction in cytoplasmic β-catenin levels caused by Hippo signalling is able to prevent the progression through the cell cycle, thus inhibiting cell proliferation.

### 3.2 Multiscale modelling of the intestinal crypt

We incorporated the subcellular Hippo/Wnt ODE formalism described above into a multiscale model of the intestinal crypt implemented in Chaste (52, 53), a computational platform previously used to simulate colonic crypt dynamics (5, 24, 25, 44, 54), amongst other multicellular systems (55, 56). The modelling of the epithelial cells that make up the intestinal crypt is organised into three interconnected modules: 1. the mechanical model, governing the cell-cell interactions and cell movement; 2. the cell-cycle model, describing how each cell progresses through the G1, S, G2 and M phases, and eventually divides; and 3. a subcellular model, describing the Hippo/Wnt signalling pathway kinetics as in the previous section.

We modelled the geometry of the crypt as a cylinder lined with epithelial cells, unrolled to a planar domain with periodic boundary conditions (Figure 2c). We coupled the cell-cycle model to the Wnt/Hippo ODE model (Supplementary Information) to account for CI effects: the YAP-P/β-catenin complex is able to cease cell-cycle progression by decreasing the level of Wnt signalling. To accomplish this, the Hippo signalling module in each cell (i.e. agent) is volume-dependent (i.e. when volume drops below a threshold the rate, 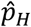 increases affecting Equations 4 and 5, see Supplementary Information for volume-dependent definition of 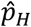). Thus, the Hippo signalling can effectively reduce the level of free β-catenin able to localise into the nucleus and promote cell-cycle progression. This mechanism causes a competition between the two pathways: an increase in Wnt shortens the cell cycle, while increasing Hippo signalling lengthens it.

We considered two different Wnt models. The first (labelled M_E_) includes Wnt as an externally-imposed gradient, and has two variants: static model (M_E1_), in which each cell reads its Wnt signal at birth (5), and dynamic model (M_E2_), in which each cell continuously updates its Wnt level, as in previous models (24, 25). In the second Wnt model (labelled M_I_), each cell contains an independent and internal level of Wnt, split with each cell division event between the two daughter cells.

### 3.3 The effects of Hippo and Wnt signalling on wild-type crypt homeostasis

#### 3.3.1 Wnt Model M_E_: externally-imposed Wnt gradient, with static (M_E1_) or dynamic (M_E2_) updating

We investigated, under physiological conditions (i.e. wild type cells), how crypt renewal dynamics are affected by CI and by the cross-talk between the Wnt (prescribed as an external gradient; models M_E1_ and M_E2_) and Hippo pathways. For this aim, we measured both the mitotic index (i.e. the percentage of cells undergoing mitosis at any given point in time) as a proxy for the proliferative capacity of cells, and the distribution of cell velocities throughout the crypt to provide a snapshot of crypt motility (Figure 3). We varied two relevant parameters: the CI threshold *V*_*thr*_ and the Wnt range (40%/100% crypt) within the crypt. The CI threshold (i.e. the cell volume below which the YAP-P/β-catenin complex prevents progression through the cell cycle, Supplementary Information) is defined as a percentage of the equilibrium cell volume enabling proliferation, and was set to either 60% or 90%; in the latter case, CI affects more cells. The Wnt level, defined as the position in the crypt at which the external Wnt gradient reaches its minimum value, is set to either 40% or 100% of the height of the crypt (Figure 2c, right panel), as an attempt to account for the proliferative differences of crypts within the small (7, 16) and large intestine (8), respectively.

**Figure 3.**
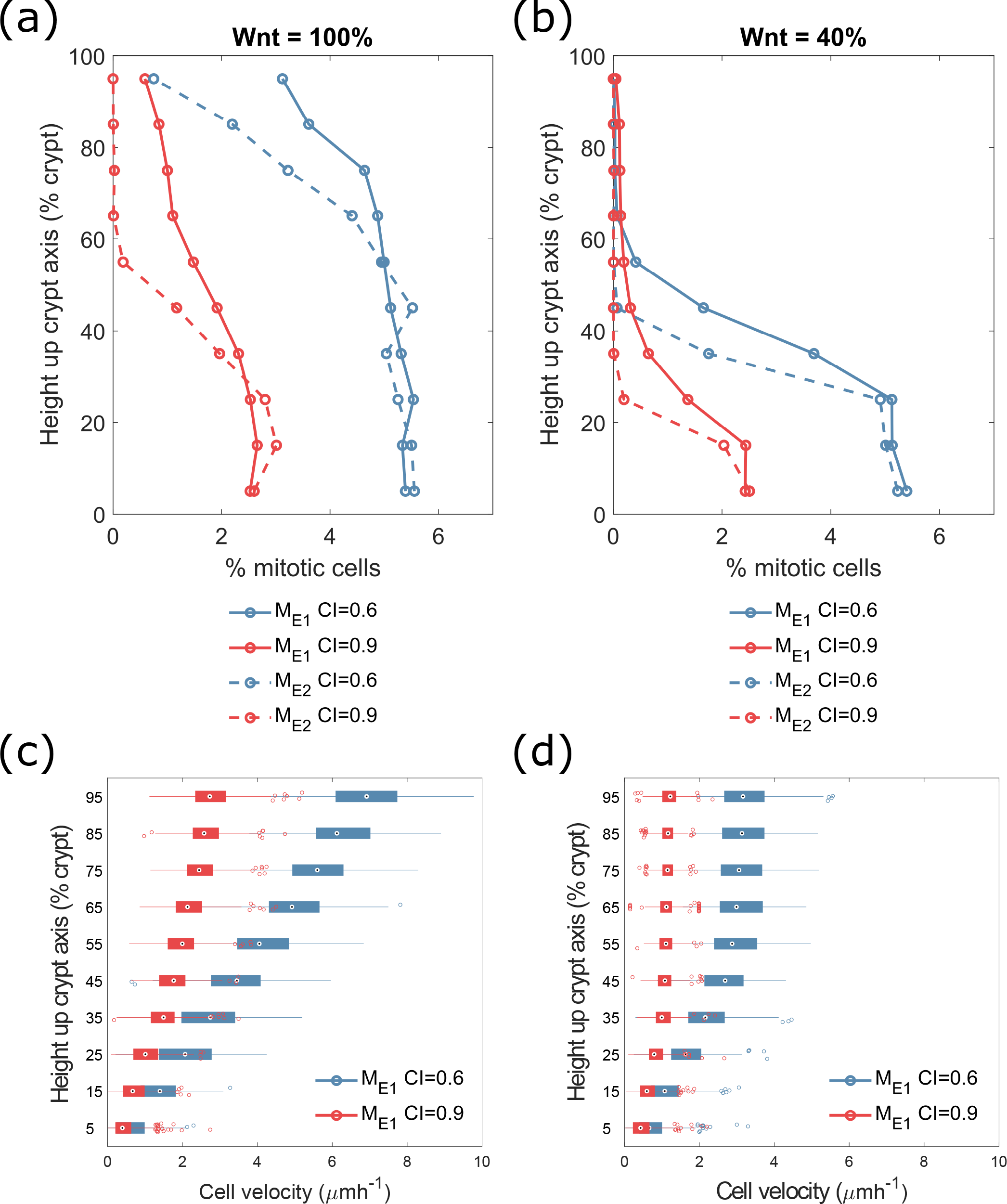
Multiscale dynamics of Hippo and Wnt signalling, and Hippo-dependent contact-inhibition (CI), in wild type crypt with imposed external Wnt gradient (M_E_) **(a, b)** Effects of Hippo-based CI on Wnt-dependent cell cycle, measuring mitotic indices (proportion of cells undergoing mitosis as a function of position in the crypt). We considered both static (M_E1_, solid lines) and dynamic (M_E2_, dashed lines) Wnt cell models (Wnt assigned at birth or continually updated, respectively), with Wnt signalling affecting 100% (a) or 40% (b) of the crypt. The different volumes at which cells undergo CI are indicated by blue (CI at 60% of equilibrium volume) and red (CI at 90% of equilibrium volume) lines. **(c, d)** Velocity whisker plots for a crypt cell population using the static (M_E1_) Wnt model, and Wnt signalling affecting 100% (c) or 40% (d) of the crypt, with CI occurring at 60% volume (blue) and 90% volume (red). The box plots indicate the median velocity of the cells at increasing heights up the crypt, with the box representing the 25^th^ and 75^th^ percentiles respectively (Supplementary Information). The whiskers extend to the most extreme velocities recorded over the course of the 150 repeated experiments.

The effect of varying Wnt level on mitotic activity (Figure 3a and b) is that, for maximal Wnt level (Figure 3a), the proliferative ‘niche’ extends further up the crypt than for lower Wnt (Figure 3b); in the latter case, there is a rapid drop-off of mitotic index at approximately 30% of the crypt height. The main difference between the static (M_E1_) and dynamic (M_E2_) Wnt models (Figures 3a and b, solid and dashed lines, respectively) is a more rapid drop-off in the mitotic index using the latter approach. This is because updating Wnt level dynamically causes cells to react instantly to changes in the external Wnt level as they move up the crypt, resulting in a more rapid cell-cycle cessation.

Varying the threshold volume at which cells are contact-inhibited (Figures 3a and b, blue and red lines corresponding to 60% and 90% of CI threshold, respectively) has a greater effect when cells reach the top of the proliferative niche where the Wnt signal decreases. This result shows that the volume-dependent prevention of β-catenin nuclear localisation, and hence also the reduction in mitotic proportion, are more pronounced when there is less free β-catenin to be sequestered. These results match our single-cell simulations, which suggested that the region over which CI is active is reduced with a lower Wnt signal. Also, our crypt simulations indicate that inhibiting the cells at 90% of their equilibrium volume results in a reduction in mitotic activity of approximately 55% at the base of the crypt and of over 75% towards the top of the proliferative ‘niche’ for both the 40% Wnt and 100% Wnt crypts. This suggests that CI within the crypt is as capable of affecting the proliferative activity of the crypt cells as changing the Wnt signalling update within the crypt.

We checked that our new model, introducing CI, does not dramatically affect the proliferative dynamics in a wild-type crypt when compared to previous models (5, 24, 25), maintaining the proliferative ‘niche’ and expected mitotic activity. Simulations (Figures 3a and b) suggest that the differential effects of Wnt on proliferation dynamics can be greater than those of moderate CI effects; still, we can tune the CI threshold to cause a comparable reduction in crypt mitotic activity.

Varying the threshold volume at which cells are contact inhibited from 60% (Figures 3c and d, blue boxes) to 90% (Figures 3c and d, red boxes) of equilibrium volume, results in a reduction in mean crypt velocity of 63%, whilst decreasing the threshold for Wnt from 100% of the crypt to 40% of the crypt (Figures 3c and d, respectively) results in a reduction in mean crypt velocity of 53%. This qualitative similarity is due to the limited effect of CI in a wild type crypt where cells are not, in general, significantly smaller than their expected equilibrium volume.

Overall, these agent-based simulations suggest Wnt signalling levels are the main regulator of cell proliferation within the crypt. Hippo signalling also plays an important role, as Hippo-dependent CI reduces proliferative activity within the crypt, with the largest effects seen in the 100% Wnt case, due to the larger number of proliferative cells affected by CI. Considering the two Wnt hypotheses for the externally-imposed Wnt model -static (M_E1_) and dynamic (M_E2_)-, the reduced mitotic activity in the M_E2_ crypt is not in line with the mitotic activity expected experimentally (6), which suggests that the static (M_E1_) modelling hypothesis is potentially more physiologically plausible; however, we would expect the effects of introduced CI to be the same in both hypotheses.

#### 3.3.2 Wnt Model M_I_: cell division-based Wnt

A prescribed and fixed external gradient of Wnt is a feature of existing computational models of crypt dynamics, effectively prescribing a spatial proliferation threshold (5, 24, 25). We modified our model to account for the aforementioned recent experimental results suggesting an alternative, division-based Wnt process (27), and investigated whether this approach could result in an emergent Wnt gradient in the crypt. We therefore set only cells at the base of the crypt to receive a maximal Wnt signal, forming a Wnt reservoir whose size (as a proportion of the height of the crypt) is a model parameter. It has been shown experimentally that Wnt does not readily diffuse into surrounding cells (27), and we therefore neglected diffusion-based spreading of Wnt. Instead, Wnt is shared stochastically between daughter cells, so that daughter cells contain a proportion 0.5±*ξ* of the mother cell’s Wnt level at each division (57, 58), where *ξ* is a sample from a Normally distributed random variable with zero mean and standard deviation σ (set as a model parameter), appropriately truncated to ensure the proportion remains in the interval [0,1].

Initial simulations, carried out with noiseless Wnt allocation on division (i.e. *σ* =0), resulted in emergent Wnt gradients over the crypt domain (Figures 4a, b). Increasing the size of the Wnt reservoir from 10% to 20% of the crypt height reduces the steepness of the resultant Wnt gradient, without affecting the mean Wnt density at the top of the crypt, so that it more closely resembles the imposed linear Wnt gradient of previous models.

**Figure 4.**
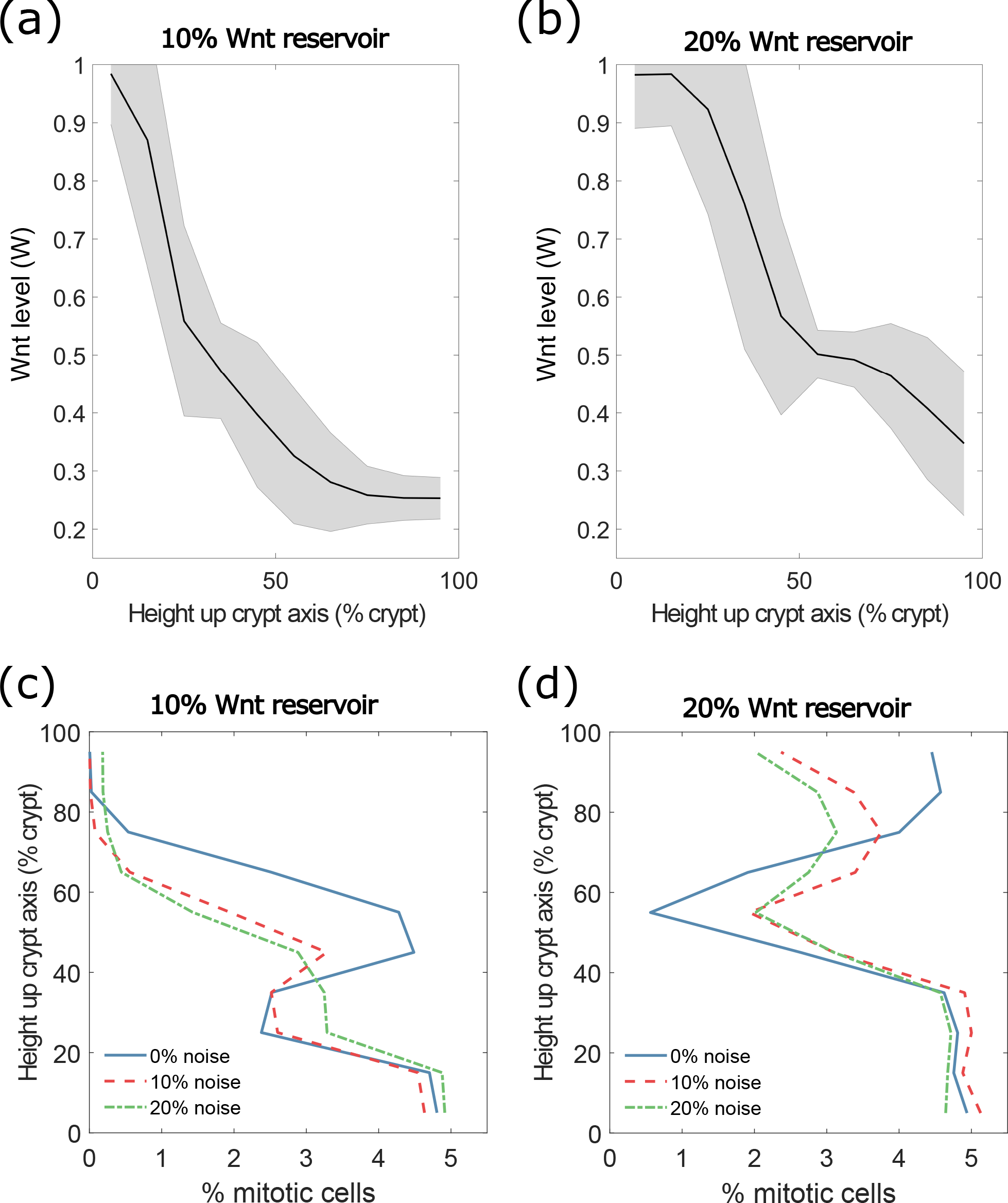
Multiscale crypt dynamics using a cell division-based Wnt model (M_I_) **(a, b)** Wnt gradient through the crypt for the cell division-based Wnt model (M_I_), with (a) 10%, and (a) 20% crypt-height Wnt reservoir. Solid line represents mean Wnt level, shaded region indicates the standard deviation amongst cells. **(c, d)** Mitotic activity, with asymmetric division of Wnt between daughter cells in crypts with (c) 10% and (d) 20% crypt-height Wnt reservoir; blue, red and green lines represent symmetric division, 10% and 20% noise, respectively.

We then introduced noise in the allocation of Wnt and repeated the wild type crypt *in silico* experiments with 10% (*σ*=0.1) and 20% (*σ*=0.2) noise, in both the 10% and 20% Wnt reservoir crypts (Figures 4c, d). The mean Wnt gradient is not strongly affected by the introduction of this division noise, as expected as the mean of the noise added is zero. Mitotic index results (Figures 4c and d) show that, considering division-based spreading of Wnt with no noise (blue lines), there is a reduction followed by a sharp increase in mitotic activity in the region where the first cell division occurs (at approximately 25%/55% of the total height of the crypt, in the case of a 10%/20% Wnt reservoir, respectively). When introducing noise in Wnt allocation upon division (Figures 4c and d, red and green lines), this reduction and subsequent increase in mitotic activity is smoother, better capturing the mitotic activity recorded experimentally (6). The latter results suggest that the distribution of Wnt between daughter cells upon cell division might not be purely symmetric, which has been suggested in a previous study (57). A comparison of externally imposed versus division-based Wnt model simulations (Figures 3a, b and Figures 4c, d, respectively) suggests that a crypt with a 10% Wnt reservoir is representative of a colonic crypt, while a 20% Wnt reservoir can resemble small intestinal crypt dynamics (6, 7). The presence of Paneth cells in the small intestinal crypt could account for this larger required Wnt reservoir that we see in our experiments; conversely, an external Wnt signal that does not extend for the full height of the crypt would also suffice in explaining this larger Wnt presence.

### 3.4 Introducing mutations in both Wnt and Hippo signalling in the crypt

Finally, we investigated the effects of CI on a dysplastic crypt; in this case, there can be a significant increase in cell number due to over-proliferation of mutant cells, resulting in a decrease in cell volume, at which point the effects of CI may play a more significant role. We considered a possible APC double-mutant (59), where there is total disruption of the β-catenin destruction complex (18, 26) as well as disruption to Hippo signalling within the cell (35); as a result, APC double-mutant cells hyper-proliferate throughout the crypt. The mutant has proliferative dynamics independent of both Wnt signal and cell volume (i.e. they are not contact inhibited). In what follows, we consider only the most realistic imposed external Wnt model (M_E1_) and the cell division-based Wnt model (M_I_). Of note, in simulating the mutant case, we used only the noise-free M_I_ model, as it has been suggested that the loss of asymmetric cell division can contribute to tumorigenesis in APC mutants (57, 58). We conducted *in silico* experiments varying the threshold volume at which Hippo signalling is active in the healthy cells, together with the size of the Wnt gradient (in M_E1_) and of the Wnt reservoir (in M_I_); Figure 5 shows mutant cell washout probabilities (i.e. the likelihood of healthy cells being able to remove the mutant from the crypt) upon insertion in the crypt of one APC double-mutant cell.

**Figure 5.**
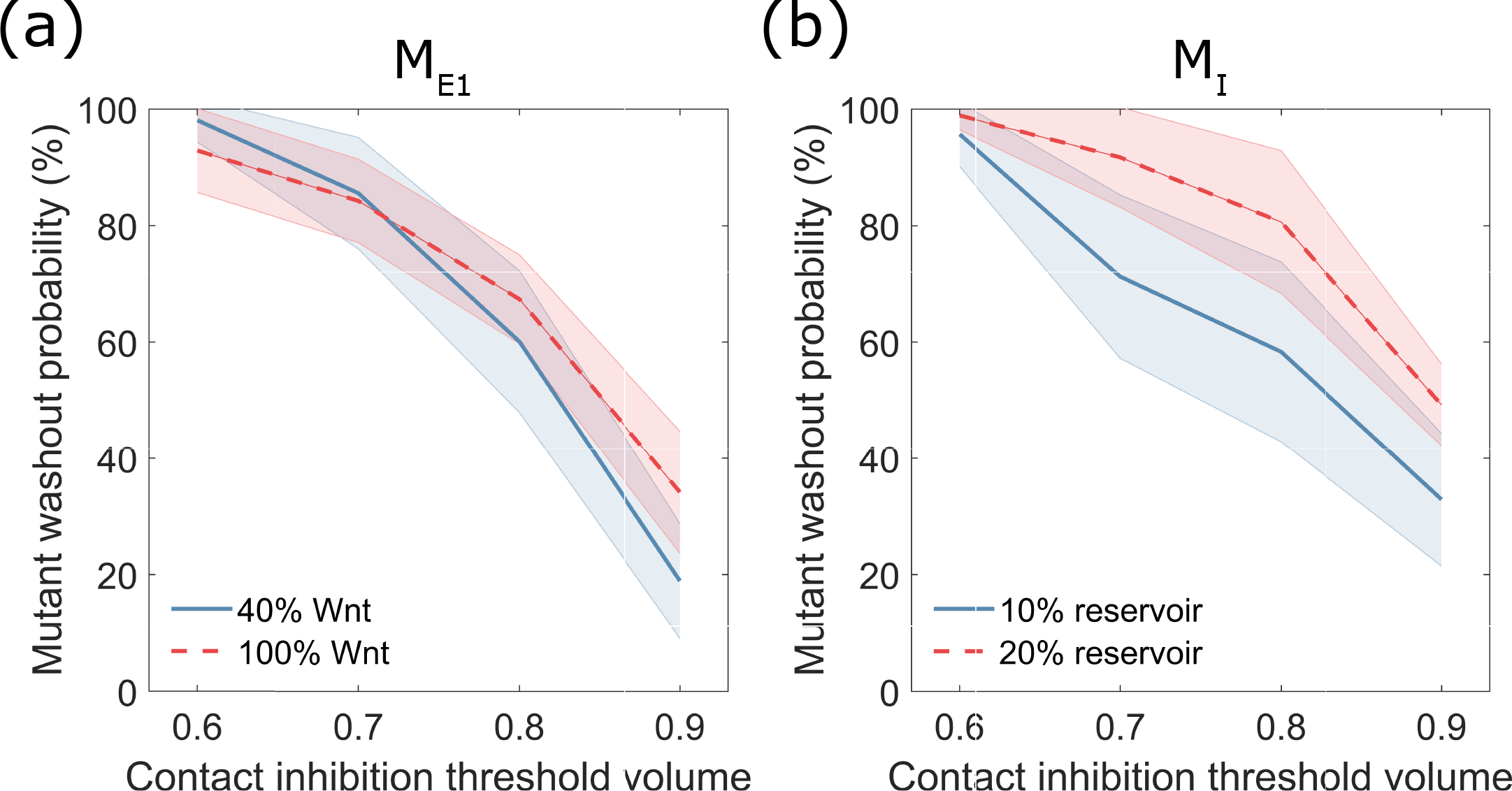
Washout probabilities upon mis-regulated Wnt and Hippo signalling activity. **(a, b)** Washout probabilities for APC double-mutant crypt, starting from a single mutant cell introduced at the base of the crypt (bottom 5% of crypt) using (a) externally-imposed gradient (M_E1_) or (b) cell-division based (M_I_) Wnt models. Blue and red lines represent the low (40% Wnt threshold/10% Wnt reservoir) and high (100% Wnt threshold/20% Wnt reservoir) Wnt cases, respectively. Increasing the threshold for CI reduces the probability of mutant washout. Shaded regions represent 95% confidence intervals for a binomial distribution with probability *p*, according to 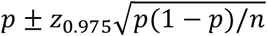, where *n* is the number of simulations conducted, and z_0.975_ is the 97.5 percentile of a standard normal distribution.

In the imposed Wnt gradient crypt (M_E1_, Figure 5a), an increase of the Wnt signal from 40% to 100% of the crypt (Figure 5a, blue and red lines, respectively) resulted in only a small change to the washout probability at low levels of CI. Conversely, at the maximal CI volume threshold (90% of equilibrium volume), the washout probability increases from approximately 20% in the 40% Wnt case to 40% in the 100% Wnt case, as the advantage of the mutant cell is reduced due to increase of proliferating healthy cells; recall that only wild type cells are affected by CI, and that mutant cells proliferate independent of cell volume and Wnt level. The increase in CI level (from 60% to 90% of equilibrium volume) reduces the washout probability by more than half in the 40% Wnt crypt (Figure 5a, blue line). It is in this case, the advantage gained by a mutant cell from the combination of its increased proliferative capacity and CI is most significant. Such reduction in washout probability is comparable with previous models, where mutant cell additional advantage was predicted to be gained through increased adhesion to the crypt substrate and to the surrounding cells (24). Of note, we did not model the altered adhesion of mutant cells, to specifically focus on signalling- and volume-dependent CI.

In the division-based Wnt model (M_I_, Figure 5b), simulations of the same mutant takeover show that, for maximal CI volume (90% of equilibrium volume), an increase in the size of the Wnt reservoir from 10% to 20% of the height of the crypt (Figure 5b, blue and red lines respectively) results in a large increase in the relative washout probability (from approximately 35% to 55%). Decreasing the volume threshold for CI resulted in a smaller decrease in the washout probability of that in the imposed-Wnt-gradient crypt with high Wnt (Figure 5, red lines), but similar behaviour for low-Wnt (Figure 5, blue lines).

Overall, the simulated dysplastic dynamics qualitatively match those predicted by the imposed Wnt gradient model, with a reduction in washout probability resulting from a stronger level of CI, and an increase in washout probability in response to a higher level of Wnt in the crypt.

## 4. Discussion

Although the role of the Wnt signalling pathway in governing the proliferative dynamics of cells within intestinal crypts has been extensively studied, a comprehensive understanding of the mechanisms linking Wnt signalling to other relevant pathways, cell mechanical properties, and cell - cycle progression in health and disease is still lacking.

In this work, we considered how two variations of a previously-proposed Wnt model affect the proliferative dynamics of intestinal crypts. We developed a new multiscale computational model of crypt dynamics, which showed that linking Wnt signalling at the point of cell division (static Wnt), as opposed to continually updating Wnt levels over the cell lifetime (dynamic Wnt), provides a more realistic representation of cell proliferation.

Previous models have assumed the existence of a fixed and externally-imposed gradient of Wnt within the crypt, with a maximal amount of Wnt found at the base, decreasing linearly with height. Based on recent experimental observations, we developed a model that assumes Wnt is internally held by each cell, supplied initially only to cells in a reservoir at the base of the crypt. We showed that subsequent division events create an emergent gradient of Wnt similar to that imposed by existing Wnt models. Noise in the Wnt allocation to daughter cells following mitosis does not significantly affect the overall Wnt gradient, but does smooth the distribution of mitotic activity within the crypt. Increasing the size of the Wnt reservoir controlled the properties of the Wnt gradient, in terms of its steepness and linearity. Furthermore, our results suggest that the spatial localisation of the Wnt reservoir can affect the crypt’s dynamics, both wild-type and dysplastic, as much as the total amount of Wnt available.

In order to elucidate the mechanisms underpinning cellular CI within the crypt, we extended a Wnt regulatory network model to additionally include a phenomenological description of Hippo signalling role in phosphorylating YAP/TAZ; the latter has been suggested to trap β-catenin within the cytoplasm and at the extracellular matrix, preventing nuclear accumulation and subsequent expression of Wnt target genes linked to cell-cycle progression (60). Our extended regulatory network is sensitive to a reduction in nuclear accumulation of β-catenin; single-cell model simulations showed that Hippo signalling can reduce nuclear β-catenin accumulation, and cause the rapid cessation of the cell cycle, matching experimental observations (35). Agent-based simulations including the Wnt/Hippo signalling coupling, with the levels of Hippo signalling further linked to cell volume, validated the single-cell analysis: the length of the cell cycle increases, and eventually proliferation ceases, as the cell volume decreases. Cells with low Wnt levels are more susceptible to CI; as the level of unphosphorylated β-catenin within the cell is reduced, the amount needed to bind to YAP-P to prevent nuclear accumulation and subsequent transcription of target genes linked to proliferation is reduced.

To investigate the effect of mutations within the crypt, we performed agent-based simulations (using both an external-gradient Wnt model with static Wnt, and the internal division-based Wnt model) with an APC double-mutant cell introduced into the crypt. The simulated mutation causes altered destruction complex kinetics in the Wnt signalling, as well as disruption of the Hippo signalling module. Increasing the amount of Wnt within the crypt did not affect the washout probability of the mutant cells at low levels of CI, with the advantage of the mutant cells not significantly reduced by the increase in proliferation caused by Wnt upregulation. The main difference arose from increasing the volume threshold for CI in the cells, with the largest decrease in the washout probability seen at a volume threshold of 90% of the healthy cell equilibrium. The effect on washout probability and cell velocity of this mutant was similar to that of increased adhesion observed previously (44). Critically, moving from the external-gradient to the internally held and division-based Wnt model shows a similar if reduced mutant advantage when implementing CI, further suggesting cell division as a plausible mechanism for the experimentally-observed Wnt gradient (27, 46). However, the greater absolute washout probability in the division-based Wnt model (~+12%) could be attributed to the shift in mitotic activity upwards in the crypt seen in this model, reducing the proliferative advantage of the mutant cell relative to the healthy cells.

The results of our study suggest the possibility of combined disease treatment, targeting both the Wnt and Hippo pathways. There are also several possible avenues for further study, to better understand the dynamics of the crypt. In the division-based Wnt model, we assumed a reservoir at the base of the crypt within which cells can uptake Wnt, as a simplification of the presence of Wnt secreting cells at the crypt base. A more detailed model should explicitly consider such cells, and Wnt secretion, in addition to possible uptake from an external reservoir. Our model also simplifies the Hippo-based sequestration of β-catenin within the cell; worthwhile extensions would be to incorporate a more detailed model of Hippo pathway components, and to explore the extent to which Hippo signalling either traps β-catenin within the cytoplasm or actively transports it to the extracellular matrix, in turn causing changes to the adhesive behaviour of the cells. Aside from Wnt and Hippo signalling, other processes also play important roles in crypt homeostasis, and would be valuable additions to a more detailed model. For example, the Notch signalling pathway coordinates cell fate specification via interaction with the Wnt pathway (61). More fundamentally, intestinal organoids represent a significant opportunity to investigate crypts *in vitro*; while computational models exist in the literature (43, 47–50), they do not all contain the level of detail described here. By introducing Wnt-secreting cells, and combining them with signalling network dependent cell-cycle models, it could be possible to create a self-sustaining organoid growth model which mimics the *in vivo* dynamics of intestinal crypts.

## Acknowledgements

LM acknowledges the support of the UK Medical Research Council (MRC, grant MR/N021444/1) and the UK Engineering and Physical Sciences Research Council (EPSRC, grant EP/R041695/1). DW was supported by an EPSRC Doctoral Training Partnership PhD studentship. AGF is supported by a Vice-Chancellor’s Fellowship from the University of Sheffield.

## Author Contribution

DW, AGF, MH and LM designed this research; DW implemented the mathematical models, generated simulations, analysed results and prepared the figures; AGF supported agent-based simulations; DW, AGF, MH and LM wrote the manuscript; MH and LM supervised the project. All authors read and approved the final manuscript.

## Competing interests

The authors have declared that no competing interests exist.

## Supplementary Information

### Single-cell modelling

The single-cell modelling we conducted combines the subcellular Wnt/Hippo signalling model, as described by a set of 9 Ordinary Differential Equations (ODEs, Figure S1) and the cell-cycle model, which is described by 5 additional ODEs (Figure S2).

The subcellular signalling model we developed is adapted from a previously published ODE model (1, 2), which describes the dynamics of β-catenin within the crypt epithelial cells, and the effects of signalling (Wnt) on the cell-cycle. The system variables, which represent protein concentrations (in [nM]), are described in the main text and in Figure 1b, with the signalling-dependent rates indicated by the coloured arrows (yellow and pink for Wnt and Hippo dependence, respectively); the parameters used for this signalling subcellular network model, and their descriptions, are included in Table S1. The derivation of the core β-catenin kinetic model can be found in the original publications (1, 2); parameters related to the adhesive, cytoplasmic and transcriptional components of β-catenin kinetics were kept as in the original model (1, 2).

The key difference with the original model is that we included a Hippo-dependent parameter to phenomenologically represent sequestration of free cytoplasmic β-catenin due to Hippo signalling. Specifically, our model describes the rate of Hippo-signalling-dependent complex (β-catenin/YAP-P, variable *C*_*H*_ in Equation (5), Figure S1); the latter can reduce the level of free β-catenin within the cell, and hence its translocation to the nucleus and subsequent transcription of cell-cycle genes (Figure 1). We described the formation of the β-catenin/YAP-P complex as being governed by Michaelis-Menten kinetics (term 1 in Equation (5), Figure S1); also, we assumed that it can localise to the cell-surface forming adhesive complexes with free cytoplasmic β-catenin (term 2 in Equations (5), Figure S1), and that it is ubiquitinated (term 3 in Equations (5), Figure S1). The β-catenin/YAP-P complex effectively links the Hippo signalling (whose strength is represented by parameter 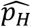 in Equation (5), Figure S1) to the level of free cytoplasmic β-catenin (*C*_*C*_, Equation (4), Figure S1), which in turn regulates the amount of transcriptionally active β-catenin (*C*_*T*_, Equation (4), Figure S1). *C*_*T*_ positively controls cell proliferation (Equation (3) in Figure S2, see description of the cell-cycle model below). Of note, 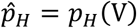 takes discrete values (and is named *p*_*H*_) in the single-cell model simulations, where it was varied from 0 to 28000h^−1^ in 500h^−1^ intervals (see section below for the definition of 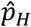 and its response to cell volume (*V*) in the agent-based simulations), and K_H_ was varied between 0nM and 20nM in 1nM intervals (see Main text and Figure 2a, b). Parameters and initial conditions for Equations (1-9) (Figure S1) are reported in Tables S1 and S2, respectively. Unlike the original model we assumed that there is only one form (open) of β-catenin and we did not consider the transcription of Wnt-target genes, which do not feedback into the other model variables.

The cell-cycle ODE model we implemented (Figure S2) was proposed in Swat et al. (4) and used in (2); it describes the interactions of 5 molecular components which control the transition of mammalian cells from the G_1_ to the S phases of the cell cycle, and includes the following proteins: phosphorylated and unphospohrylated retinoblastoma proteins (*pRB*_*p*_ and *pRB*, respectively), the transcription factor E2F1, and active and inactive cyclin D kinase complexes (*CycD*_*a*_ and *CycD*_*i*_ respectively). The level of E2F1 has been shown to positively regulate the G_1_ to S phase transition (5). In the equations in Figure S2 (1-5), parameters *k* and *ϕ* are non-negative rate constants, parameters *K* and *J* are Michaelis-Menten dissociation constants, and *C*_*T*_ is the level of transcriptional β-catenin, as governed by Equation (9) in Figure S1. Thus, the interplay between crypt signalling (Wnt/Hippo) and the cell-cycle is governed by the level of transcriptional β-catenin (*C*_*T*_), as CycD_i_ levels (Equation (3), Figure S2) depend on Wnt signalling. As the level of Wnt signalling is decreased, the production of C_T_ is reduced; this results in a decrease of both CycD expression, and of the downstream cell-cycle effector E2F1, preventing cell transitioning from the G_1_ to the S cell-cycle phase. The parameters and initial conditions for the cell-cycle model remained set as in Swat et al. (4); parameters reflect experimentally known features of each molecular component involved in the cell-cycle, with the fitting procedure described in the original publication.

The combined subcellular representation of the Wnt/Hippo signalling and the cell-cycle is therefore comprised of 14 ODEs (combined Equations in Figure S1 and S2). The ODEs were solved using the ODE45 function in MATLAB R2017b, with a timestep of 0.005h, for a total simulation duration of 40h. The length of the G_1_ phase of the cell cycle (Figure 2 a, b) was determined by the time taken for the concentration of E2F1 in the cell to exceed 1, as identified in Swat et al. (4).

### Agent-based modelling

Our agent-based model was implemented within the Chaste (v. 3.3) modelling framework (6, 7). Chaste is set-up to allow for modular model compositions that span multiple spatial scales; it incorporates features for cellular level dynamics, in the form of the mechanical cell modelling, as well as subcellular features such as gene-regulatory networks or alternative subcellular models, and provides tools for incorporating cell-cycle models that can combine with the subcellular model for multi-scale description of cell population dynamics.

The model for the mechanical interactions between the cells uses a cell-centred design, under the assumption that the cells within the epithelial sheet that forms the surface of the intestinal crypt are adhered to one another (8); in such a way, cells provide forces that push neighbouring cells away, and no holes in the epithelial layer are present. Each cell is modelled as a single point, the centre of the cell, connected to each of its neighbours by a spring, as in (2). Cell boundaries are formed by completing a Voronoi tessellation across the surface. The volume of each cell is also calculated using this Voronoi tessellation, and determines the level of Hippo signalling active in the cell, as described below.

The subcellular Wnt/Hippo signalling and the cell-cycle model, and related parameters, remained as in the single-cell model. The only differences are the rate 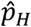 and the constant *K*_*H*_ (Equation (5) Figure S1). The rate 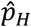 is now a function of volume described as follows:

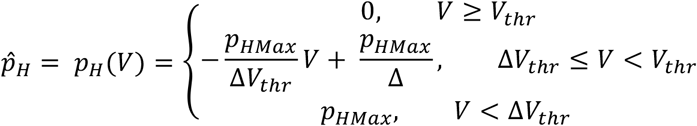

where *V*_*thr*_ is the threshold volume for the onset of contact inhibition, *p*_*HMax*_ is the maximal rate of Hippo dependent β-catenin sequestration (set using the single cell model as shown in Figure 2), *V*_*thr*_ is the volume threshold at which Hippo is active, and Δ describes the volume reduction required to cause complete cell-cycle cessation. The value for Δ is fitted to allow cell-cycle length elongation before cessation, with Δ = 0 effectively causing contact inhibition to occur instantaneously when the cell volume drops below the threshold *V*_*thr*_. Including Δ > 0 maintains the proliferative region towards the crypt base. The process of cell-cycle cessation due to volume reduction has been previously implemented as a switch (9), stopping cell-cycle progression below a set volume, while our implementation results in smoother transitions. The value of *V*_*thr*_ is varied in our agent-based simulations between 0.6 and 0.9, and *p*_*HMax*_ remains fitted as in the single cell analysis (Table S1).

In the agent-based model, ODEs were solved using the Sundials CVODE solving tools, which dynamically varies the timestep and solving method depending on the system.

The crypt domain was modelled as an unrolled cylinder, allowing for the projection of cells from the three-dimensional physiological space to a two-dimensional modelling space, containing approximately 14 cells in diameter and 19 cells in high, as suggested by mouse experimental data (10). The unrolled cylinder representation has periodic boundaries in the X domain, a solid boundary at Y = 0 and a sloughing boundary at the top of the crypt (Y=Y_max_) which removes those cells whose cell centres pass it. Our equilibrium cell volume, calculated as volume of proliferative cells, was set as 0.74, taken from the volume distribution of a relaxed crypt. An artefact of the 2D cylindrical representation of the cells is that those cells on the base boundary (Y=0) appear significantly smaller, which causes these cells to be more contact-inhibited. To account for this, cells with volumes below 0.15 (calculated from the distribution of equilibrium crypt volumes) were scaled by a factor of 5, such that the volume distribution at the base could match the volume distribution in the rest of the proliferative region of the crypt. The threshold volume for contact inhibition (*V*_*c*_) is a variable parameter in our investigation (see Main Text).

In agent-based simulations, we coupled the cell-cycle model to the level of transcription complexes *C*_*T*_ (Figure 1b), as in (2) and in the single-cell model described above. The greater Wnt signal at the base of the crypt causes the cell to produce Cyclin D and thus progress through the cell cycle more rapidly, ultimately creating an increasing age gradient over the crypt, and the appearance of a proliferative base of the crypt, as seen experimentally (10). This cell-cycle model was applied to both the externally-imposed gradient (M_E_) and division-based (M_I_) Wnt models.

Agent-based simulations (Figures 3-5) were carried out with a timestep of 0.01h; before the *in silico* experiments were started, the crypt was allowed to reach equilibrium by simulating an initial 300 hours to allow cell volumes to relax from the initial honeycomb tessellation and the subcellular model to reach equilibrium. Each experiment, of 1000 (simulated) hours duration, was repeated 150 times to provide sufficient data for statistical analysis.

The box plots in Figure 3c, d show the median velocity of the cells at increasing heights up the crypt, with the box representing the 25^th^ and 75^th^ percentiles respectively. The whiskers extend to the most extreme velocities recorded over the course of the 150 repeated experiments. The error bars in Figure 4 represent the mean and standard deviation across the repeated experiments. Mitotic proportions (Figures 3, 4) are defined as the proportion of cells residing in the M phase of the cell cycle, averaged over time, at a given point in the crypt (mean over a region of size 10% of the total crypt height). Mitotic proportion is thus a measure of the proliferative capacity of the cells, as it indicates that cells are progressing through the cell cycle.

The shaded regions in Figure 5 denote the 95% confidence interval for a binomial distribution with probability *p*, according to 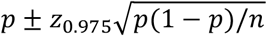, where *n* is the number of simulations conducted, and z_0.975_ is the 97.5 percentile of a standard normal distribution. In the mutant cell simulations, one mutant cell was introduced at the base of the crypt (in a random location along the first row of cells) at the start of the simulation; the washout probability was defined as the proportion of simulations where there were no mutant cells left within the crypt at the end of the experiment (after 1000 hours).

**Figure S1.**
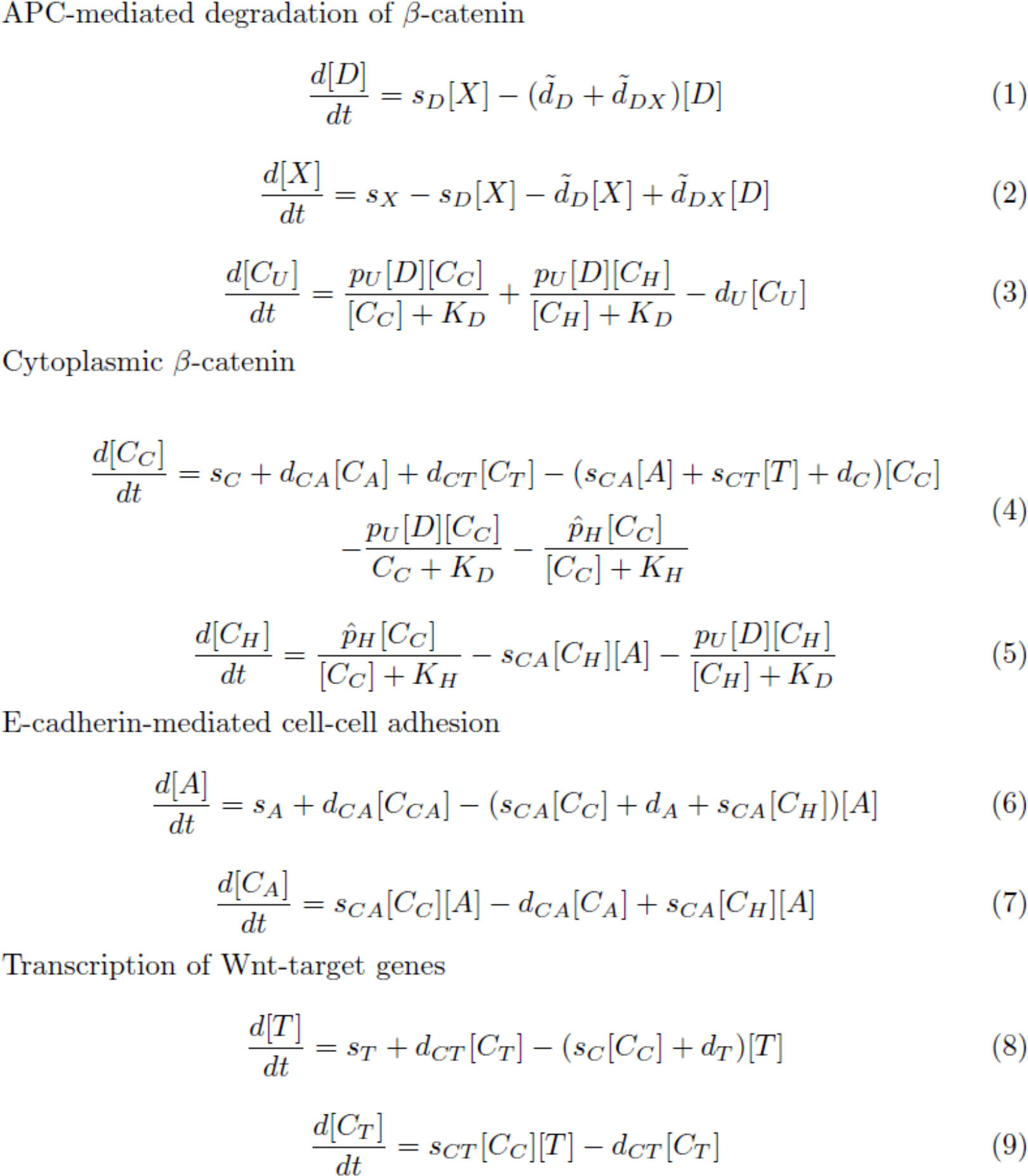
Ordinary differential equations of the kinetic model of β-catenin localisation shown in Figure 1, describing the change of β-catenin complex concentrations and their link to cell-cycle progression. *C*_*A*_, *C*_*C*_, *C*_*T*_, and *C*_*U*_ are the levels of adhesive-linked β-catenin at the cell surface, cytosolic β-catenin, transcriptional nuclear β-catenin, and β-catenin marked for degradation, respectively. *A*, *T*, and *D* denote the concentrations of molecular species forming complexes with β-catenin at the cell surface (forming adhesive complexes at the adherens junction), within the nucleus, and within the destruction complex, respectively. *X* denotes the concentration of Axin. *C*_*H*_ denotes the concentration of β-catenin/YAP complex formed due to Hippo signalling in the cell. The symbols carrying tildes (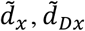 and 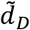) are those that may vary in response to Wnt signalling. It was assumed that these parameters are linear functions of Wnt signal (W), which varies from W = 0 to W = 1 within the prescribed Wnt region (40%/100%). They are described as follows: 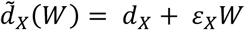, 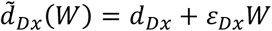 and 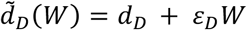. Parameter are described above, and their values are reported in Table S1.

**Figure S2.**
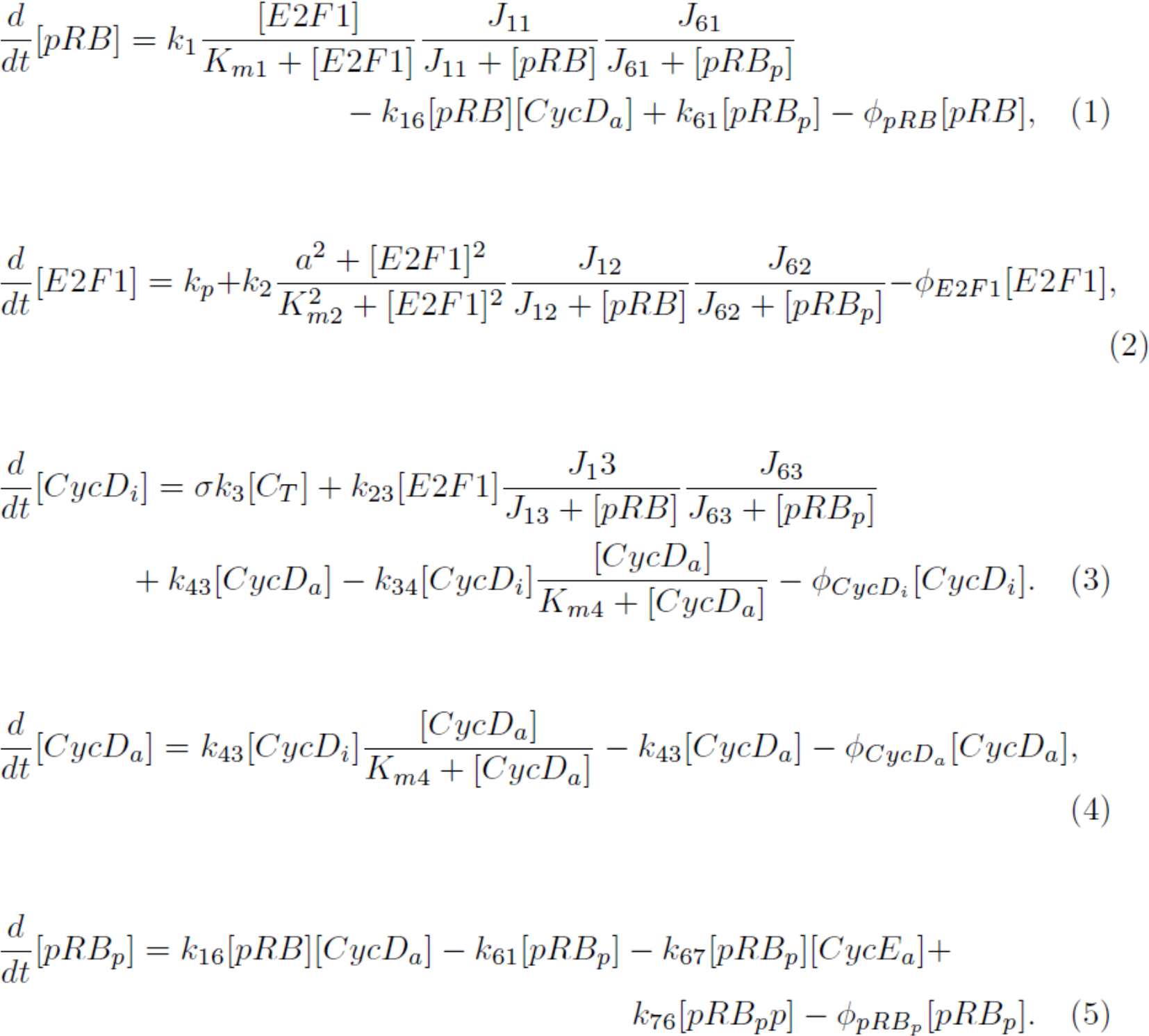
Ordinary differential equations governing cell-cycle progression, linked to the kinetic model of β-catenin localisation shown in Figure 1 (ODEs in Equations S1) through the levels of *C*_*T*_ which affect CycD level. The model includes 5 molecular components which control the transition of mammalian cells from the G_1_ to the S phases of the cell cycle: phosphorylated and unphospohrylated retinoblastoma proteins (*pRB*_*p*_ and *pRB*, respectively), the transcription factor E2F1, active and inactive cyclin D kinase complexes (*CycD*_*a*_ and *CycD*_*i*_ respectively). To link the subcellular β-catenin model with the cell-cycle model we apply a mitogenic factor, *σ* = 1/25 (Equation 3). Further details and parameter values can be found in the original papers (2, 4).

**Table S1.**
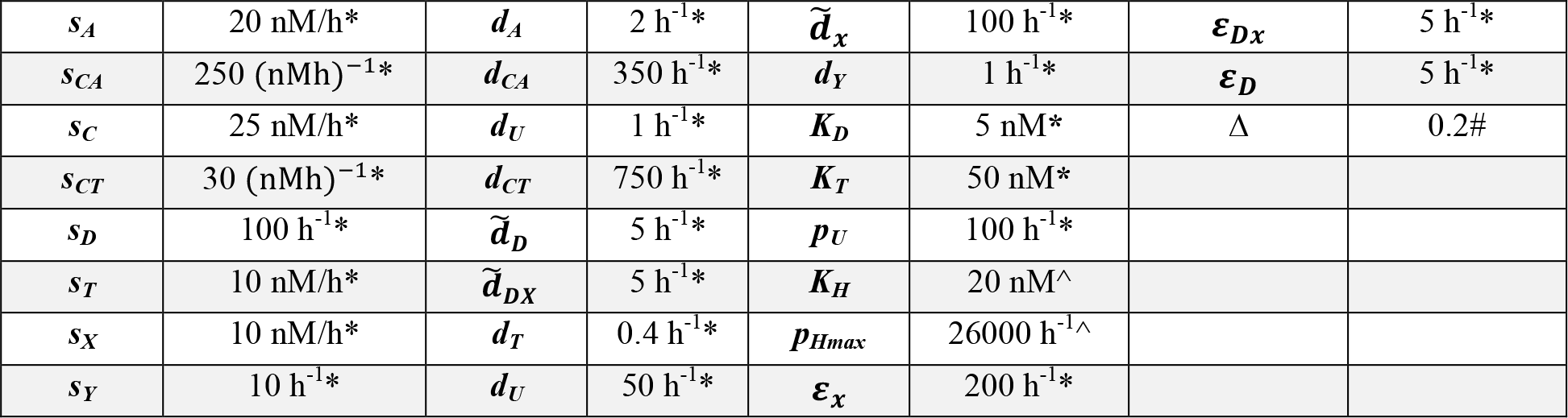
Table of model parameter values (ODEs in Figure S1). *s*_*A*_ is the basal rate of adhesive molecule formation; *s*_*CA*_ is the formation rate of adhesive β-catenin complexes; *s*_*c*_ is the rate of production of β-catenin within the cytosol; *s*_*CT*_ is the formation rate of transcriptional β-catenin; *s*_*D*_ is the rate of formation of the destruction complex; *s*_*T*_ is the basal rate of transcriptional molecule production; *s*_*X*_ is the basal rate of Axin production; *s*_*Y*_ is the maximal rate of transcription of Wnt target genes; *d*_*A*_ is the degradation rate for adhesive molecules; *d*_*CA*_ is the disassociation rate of adhesive complexes; *d*_*C*_ is the APC independent degradation rate of β-catenin; *d*_*CT*_ is the dissociation rate of transcriptional complexes; 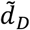 is the Wnt dependent degradation of the destruction complex (linearly dependent on Wnt according to the equations in the legend of Figure S1); 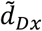 is the Wnt dependent disassociation rate for the destruction complex into Axin; *d*_*T*_ is the degradation rate for transcriptional molecules; *d*_*CU*_ is the degradation rate of ubiquitinated β-catenin; 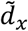 is the Wnt dependent degradation rate of Axin; *d*_*Y*_ is the rate of transcription of Wnt target proteins; *K*_*D*_ is a saturation term for the ubiquitination of β-catenin by the destruction complex; *K*_*T*_ is the saturation term for the transcription of Wnt target proteins due to the nuclear accumulation of β-catenin; *p*_*U*_ is the association term form the destruction complex and cytoplasmic β-catenin for ubiquitination; *K*_*H*_ is the saturation term for the formation of a YAP-P/β-catenin complex; 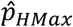 is the Hippo (hence volume) dependent association rate for YAP-P and β-catenin. Δ describes the volume reduction required to cause complete cell-cycle cessation.* parameters as in (1, 2), ^ parameters fitted in single-cell analysis, # parameter estimated using agent-based modelling, such that the reduction in volume causes cell cycle length elongation before cessation, maintaining the proliferative niche towards the crypt base.

|  |  |  |  |  |  |  |  |
| --- | --- | --- | --- | --- | --- | --- | --- |
| $s_A$ | 20 nM/h* | $d_A$ | 2 h<sup>-1</sup>* | $\tilde{d}_x$ | 100 h<sup>-1</sup>* | $\epsilon_{Dx}$ | 5 h<sup>-1</sup>* |
| $s_{CA}$ | 250 (nMh)<sup>-1</sup>* | $d_{CA}$ | 350 h<sup>-1</sup>* | $d_Y$ | 1 h<sup>-1</sup>* | $\epsilon_D$ | 5 h<sup>-1</sup>* |
| $s_C$ | 25 nM/h* | $d_U$ | 1 h<sup>-1</sup>* | $K_D$ | 5 nM* | $\Delta$ | 0.2# |
| $s_{CT}$ | 30 (nMh)<sup>-1</sup>* | $d_{CT}$ | 750 h<sup>-1</sup>* | $K_T$ | 50 nM* | | |
| $s_D$ | 100 h<sup>-1</sup>* | $\tilde{d}_D$ | 5 h<sup>-1</sup>* | $p_U$ | 100 h<sup>-1</sup>* | | |
| $s_T$ | 10 nM/h* | $\tilde{d}_{Dx}$ | 5 h<sup>-1</sup>* | $K_H$ | 20 nM<sup>^</sup> | | |
| $s_X$ | 10 nM/h* | $d_T$ | 0.4 h<sup>-1</sup>* | $p_{Hmax}$ | 26000 h<sup>-1</sup><sup>^</sup> | | |
| $s_Y$ | 10 h<sup>-1</sup>* | $d_U$ | 50 h<sup>-1</sup>* | $\epsilon_x$ | 200 h<sup>-1</sup>* | | |

**Table S2.** Table of initial conditions for subcellular Wnt model described in Figure S1.

|  |  |  |  |  |  |
| --- | --- | --- | --- | --- | --- |
| [A*] | 10 nM | [C<sub>T</sub>] | 2.54 nM | [X] | 0.067 nM |
| [C<sub>A</sub>*] | 18.14 nM | [D] | 0.67 nM | [C<sub>U</sub>] | 0.45 nM |
| [C<sub>C</sub>*] | 2.54 nM | [T] | 25 nM | [C<sub>H</sub>] | 0.0 nM |

## References

1. Ferlay J, Soerjomataram I, Dikshit R, Eser S, Mathers C, Rebelo M, et al. Cancer incidence and mortality worldwide: Sources, methods and major patterns in GLOBOCAN 2012. International Journal of Cancer. 2015.

2. Bosetti C, Levi F, Rosato V, Bertuccio P, Lucchini F, Negri E, et al. Recent trends in colorectal cancer mortality in Europe. Int J Cancer. 2011;129(1):180–91.

3. Young M, Ordonez L, Clarke AR. What are the best routes to effectively model human colorectal cancer? Mol Oncol. 2013;7(2):178–89.

4. Baker AM, Cereser B, Melton S, Fletcher AG, Rodriguez-Justo M, Tadrous PJ, et al. Quantification of crypt and stem cell evolution in the normal and neoplastic human colon. Cell Reports. 2014;8(4):940–7.

5. Dunn S-J, Osborne JM, Appleton PL, Näthke I. Combined changes in Wnt signaling response and contact inhibition induce altered proliferation in radiation-treated intestinal crypts. Molecular Biology of the Cell. 2016;27:1863–74.

6. Trani D, Nelson SA, Moon B-H, Swedlow JJ, Williams EM, Strawn SJ, et al. High-Energy Particle-Induced Tumorigenesis Throughout the Gastrointestinal Tract. Radiation Research. 2014;181(2):162–71.

7. Potten CS, Kellett M, Rew DA, Roberts SA. Proliferation in human gastrointesinal epithelium using bromodeoxyuridine in vivo: Data for different sites, proximity to a tumour, and polyposis coli. Gut. 1992;33(4):524–9.

8. Sunter JP, Appleton DR, Dé Rodriguez MS, Wright Na, Watson AJ, De Rodriguez MS. A comparison of cell proliferation at different sites within the large bowel of the mouse. Journal of anatomy. 1979;129:833–42.

9. Stamos JL, Weis WI. The β-catenin destruction complex. Cold Spring Harbor Perspectives in Biology. 2013;5(1).

10. Reya T, Clevers H. Wnt signalling in stem cells and cancer. Nature. 2005;434(7035):843–50.

11. Ilyas M. Wnt signalling and the mechanistic basis of tumour development. Journal of Pathology. 2005;205(2):130–44.

12. Lloyd-Lewis B, Fletcher AG, Dale TC, Byrne HM. Toward a quantitative understanding of the Wnt/β-catenin pathway through simulation and experiment. Wiley Interdisciplinary Reviews: Systems Biology and Medicine. 2013;5(4):391–407.

13. Sumigray KD, Terwilliger M, Lechler T. Morphogenesis and Compartmentalization of the Intestinal Crypt. Developmental Cell. 2018;45(2):183–97.

14. Van de Wetering M, Sancho E, Verweij C, De Lau W, Oving I, Hurlstone A, et al. The β-catenin/TCF-4 complex imposes a crypt progenitor phenotype on colorectal cancer cells. Cell. 2002;111:241–50.

15. Clevers H, Nusse R. Wnt/β-catenin signaling and disease. Cell. 2012;149(6):1192–205.

16. Potten CS, Kellett M, Roberts SA, Rew DA, Wilson GD. Measurement of in vivo proliferation in human colorectal mucosa using bromodeoxyuridine. Gut. 1992;33(1):71–8.

17. Kwong LN, Dove WF. APC and its modifiers in colon cancer. Advances in experimental medicine and biology. 2009;656:85–106.

18. Lamlum H, Papadopoulou A, Ilyas M, Rowan A, Gillet C, Hanby A, et al. APC mutations are sufficient for the growth of early colorectal adenomas. Proceedings of the National Academy of Sciences. 2000;97(5):2225–8.

19. Fre S, Huyghe M, Mourikis P, Robine S, Louvard D, Artavanis-Tsakonas S. Notch signals control the fate of immature progenitor cells in the intestine. Nature. 2005;435:964–8.

20. Wright NAA, Malcolm. The biology of epithelial cell populations. Oxford: Oxford University Press; 1984.

21. Varelas X, Miller BW, Sopko R, Song S, Gregorieff A, Fellouse FA, et al. The Hippo Pathway Regulates Wnt/β-Catenin Signaling. Developmental Cell. 2010;18:579–91.

22. Swat M, Kel A, Herzel H. Bifurcation analysis of the regulatory modules of the mammalian G 1 / S transition. Bioinformatics. 2004;20:1506–11.

23. Scoville DH, Sato T, He XC, Li L. Current View: Intestinal Stem Cells and Signaling. Gastroenterology. 2008;134(3):849–64.

24. Mirams GR, Fletcher AG, Maini PK, Byrne HM. A theoretical investigation of the effect of proliferation and adhesion on monoclonal conversion in the colonic crypt. Journal of Theoretical Biology. 2012;312:143–56.

25. van Leeuwen IMM, Mirams GR, Walter A, Fletcher AG, Murray PJ, Osborne J, et al. An integrative computational model for intestinal tissue renewal. Cell Proliferation. 2009;42:617–36.

26. Ingham-Dempster T, Corfe B, Walker D. A cellular based model of the colon crypt suggests novel effects for Apc phenotype in colorectal carcinogenesis. J Comput Sci-Neth. 2018;24:125–31.

27. Farin HF, Jordens I, Mosa MH, Basak O, Korving J, Tauriello DV, et al. Visualization of a short-range Wnt gradient in the intestinal stem-cell niche. Nature. 2016;530(7590):340–3.

28. Dong J, Feldmann G, Huang J, Wu S, Zhang N, Comerford SA, et al. Elucidation of a Universal Size-Control Mechanism in Drosophila and Mammals. Cell. 2007;130:1120–33.

29. Helmlinger G, Netti PA, Lichtenbeld HC, Melder RJ, Jain RK. Solid stress inhibits the growth of multicellular tumor spheroids. Nature Biotechnology. 1997;15:778–83.

30. Eisenhoffer GT, Loftus PD, Yoshigi M, Otsuna H, Chien CB, Morcos PA, et al. Crowding induces live cell extrusion to maintain homeostatic cell numbers in epithelia. Nature. 2012;484(7395):546–9.

31. Zeng Q, Hong W. The Emerging Role of the Hippo Pathway in Cell Contact Inhibition, Organ Size Control, and Cancer Development in Mammals. Cancer Cell. 2008;13(3):188–92.

32. Zhao B, Zhao B, Wei X, Wei X, Li W, Li W, et al. Inactivation of YAP oncoprotein by the Hippo pathway is involved in cell contact inhibition and tissue growth control. Genes & development. 2007;21:2747–61.

33. Camargo FD, Gokhale S, Johnnidis JB, Fu D, Bell GW, Jaenisch R, et al. YAP1 Increases Organ Size and Expands Undifferentiated Progenitor Cells. Current Biology. 2007;17:2054–60.

34. Barry ER, Camargo FD. The Hippo superhighway: Signaling crossroads converging on the Hippo/Yap pathway in stem cells and development. Current Opinion in Cell Biology. 2013;25:247–53.

35. Imajo M, Miyatake K, Iimura A, Miyamoto A, Nishida E. A molecular mechanism that links Hippo signalling to the inhibition of Wnt/β-catenin signalling. EMBO Journal. 2012;31:1109–22.

36. Dupont S, Morsut L, Aragona M, Enzo E, Giulitti S, Cordenonsi M, et al. Role of YAP/TAZ in mechanotransduction. Nature. 2011;474:179–84.

37. Jaalouk DE, Lammerding J. Mechanotransduction gone awry. Nature Reviews Molecular Cell Biology. 2009;10(1):63–73.

38. Johnson R, Halder G. The two faces of Hippo: targeting the Hippo pathway for regenerative medicine and cancer treatment. Nature Reviews Drug Discovery. 2013;13:63–79.

39. Yang C-C, Graves HK, Moya IM, Tao C, Hamaratoglu F, Gladden AB, et al. Differential regulation of the Hippo pathway by adherens junctions and apical–basal cell polarity modules. Proceedings of the National Academy of Sciences. 2015;112:1785–90.

40. Schroeder MC, Halder G. Regulation of the Hippo pathway by cell architecture and mechanical signals. Seminars in Cell and Developmental Biology. 2012;23(7):803–11.

41. Piccolo S, Dupont S, Cordenonsi M. The Biology of YAP/TAZ: Hippo Signaling and Beyond. Physiological Reviews. 2014;94:1287–312.

42. van Leeuwen IMM, Byrne HM, Jensen OE, King JR. Elucidating the interactions between the adhesive and transcriptional functions of β-catenin in normal and cancerous cells. Journal of Theoretical Biology. 2007;247:77–102.

43. Fletcher AGM, Phillip.; Maini, Phillip. Multiscale modelling of intestinal crypt organization and carcinogenesis. Mathematical Models and Methods in Applied Sciences. 2015;484:546–9.

44. Fletcher AG, Breward CJW, Jonathan Chapman S. Mathematical modeling of monoclonal conversion in the colonic crypt. Journal of Theoretical Biology. 2012;300:118–33.

45. Farin HF, Van Es JH, Clevers H. Redundant sources of Wnt regulate intestinal stem cells and promote formation of paneth cells. Gastroenterology. 2012;143(6):1518–29.

46. Shoshkes-Carmel M, Wang YJ, Wangensteen KJ, Tóth B, Kondo A, Massassa EE, et al. Subepithelial telocytes are an important source of Wnts that supports intestinal crypts. Nature. 2018;557(7704):242–6.

47. Lee E, Salic A, Krüger R, Heinrich R, Kirschner MW. The roles of APC and axin derived from experimental and theoretical analysis of the Wnt pathway. PLoS Biology. 2003;1(1):116–32.

48. Mirams GR, Byrne HM, King JR. A multiple timescale analysis of a mathematical model of the Wnt/beta-catenin signalling pathway. Journal of mathematical biology. 2010;60(1):131–60.

49. Schmitz Y, Rateitschak K, Wolkenhauer O. Analysing the impact of nucleo-cytoplasmic shuttling of β-catenin and its antagonists APC, Axin and GSK3 on Wnt/β-catenin signalling. Cellular Signalling. 2013;25(11):2210–21.

50. MacLean AL, Rosen Z, Byrne HM, Harrington HA. Parameter-free methods distinguish Wnt pathway models and guide design of experiments. Proceedings of the National Academy of Sciences. 2015;112(9):2652–7.

51. Pedone E, Olteanu V-A, Marucci L, Muñoz-Martin MI, Youssef SA, de Bruin A, et al. Modeling Dynamics and Function of Bone Marrow Cells in Mouse Liver Regeneration. Cell Reports. 2017;18(1):107–21.

52. Pitt-Francis J, Pathmanathan P, Bernabeu MO, Bordas R, Cooper J, Fletcher AG, et al. Chaste: A test-driven approach to software development for biological modelling. Computer Physics Communications. 2009;180:2452–71.

53. Mirams GR, Arthurs CJ, Bernabeu MO, Bordas R, Cooper J, Corrias A, et al. Chaste: An Open Source C++ Library for Computational Physiology and Biology. PLoS Computational Biology. 2013;9.

54. Dunn SJ, Fletcher AG, Chapman SJ, Gavaghan DJ, Osborne JM. Modelling the role of the basement membrane beneath a growing epithelial monolayer. Journal of Theoretical Biology. 2012;298:82–91.

55. Osborne JM, Fletcher AG, Pitt-Francis JM, Maini PK, Gavaghan DJ. Comparing individual-based approaches to modelling the self-organization of multicellular tissues. PLoS Computational Biology. 2017;13(2):e1005387.

56. Godwin S, Ward D, Pedone E, Homer M, Fletcher AG, Marucci L. An extended model for culture-dependent heterogenous gene expression and proliferation dynamics in mouse embryonic stem cells. NPJ Syst Biol Appl. 2017;3:19.

57. Qu Y, Gharbi N, Yuan X, Olsen JR, Blicher P, Dalhus B, et al. Axitinib blocks Wnt/β-catenin signaling and directs asymmetric cell division in cancer. Proceedings of the National Academy of Sciences. 2016;113(33):9339–44.

58. Lien WH, Fuchs E. Wnt some lose some: transcriptional governance of stem cells by Wnt/beta-catenin signaling. Genes & Development. 2014;28(14):1517–32.

59. Cai J, Maitra A, Anders Ra, Taketo MM, Pan D. β-Catenin destruction complex-independent regulation of Hippo – YAP signaling by APC in intestinal tumorigenesis. Genes & development. 2015:1–14.

60. Miyaoka Y, Miyajima A. To divide or not to divide: Revisiting liver regeneration. Cell Division. 2013;8(1):8.

61. Kay SK, Harrington HA, Shepherd S, Brennan K, Dale T, Osborne JM, et al. The role of the Hes1 crosstalk hub in Notch-Wnt interactions of the intestinal crypt. PLoS Computational Biology. 2017;13(2):e1005400.

## References

1. van Leeuwen IMM, Byrne HM, Jensen OE, King JR. Elucidating the interactions between the adhesive and transcriptional functions of β-catenin in normal and cancerous cells. Journal of Theoretical Biology. 2007;247:77–102.

2. van Leeuwen IMM, Mirams GR, Walter A, Fletcher AG, Murray PJ, Osborne J, et al. An integrative computational model for intestinal tissue renewal. Cell Proliferation. 2009;42:617–36.

3. Imajo M, Miyatake K, Iimura A, Miyamoto A, Nishida E. A molecular mechanism that links Hippo signalling to the inhibition of Wnt/β-catenin signalling. EMBO Journal. 2012;31:1109–22.

4. Swat M, Kel A, Herzel H. Bifurcation analysis of the regulatory modules of the mammalian G 1 / S transition. Bioinformatics. 2004;20:1506–11.

5. Kel A, Deineko I, Kel-Margoulis O, Wingender E, Ratner V. Modelling of gene regulatory networks of cell cycle control. Role of E2F feedback loops. German conference on Bioinformatics GCB. 2000:107–114

6. Pitt-Francis J, Pathmanathan P, Bernabeu MO, Bordas R, Cooper J, Fletcher AG, et al. Chaste: A test-driven approach to software development for biological modelling. Computer Physics Communications. 2009;180:2452–71.

7. Mirams GR, Arthurs CJ, Bernabeu MO, Bordas R, Cooper J, Corrias A, et al. Chaste: An Open Source C++ Library for Computational Physiology and Biology. PLoS Computational Biology. 2013;9.

8. Meineke FA, Potten CS, Loeffler M. Cell migration and organization in the intestinal crypt using a lattice-free model. Cell Proliferation. 2001;34:253–66.

9. Dunn S, Osborne J, Appleton P, Näthke I. Combined changes in Wnt signaling response and contact inhibition induce altered proliferation in radiation-treated intestinal crypts. Molecular Biology Of The Cell. 2016; 27(11), 1863–1874.

10. Sunter JP, Appleton DR, Dé Rodriguez MS, Wright Na, Watson AJ, De Rodriguez MS. A comparison of cell proliferation at different sites within the large bowel of the mouse. Journal of anatomy. 1979;129:833–42.

